# Chikungunya virus transmission in the Southernmost state of Brazil was characterized by self-limited cases (2017–2019) and a larger 2021 outbreak

**DOI:** 10.1101/2022.09.30.510389

**Authors:** Tatiana Schäffer Gregianini, Richard Steiner Salvato, Regina Bones Barcellos, Fernanda Marques Godinho, Amanda Pellenz Ruivo, Viviane Horn de Melo, Júlio Augusto Schroder, Fernanda Letícia Martiny, Erica Bortoli Möllmann, Cátia Favreto, Ludmila Fiorenzano Baethgen, Vithoria Pompermaier Ferreira, Lívia Eidt de Lima, Cláudia Fasolo Piazza, Taís Raquel Marcon Machado, Irina Marieta Becker, Raquel Rocha Ramos, Guilherme Carey Frölich, Alana Fraga Rossetti, Lucas da Cunha Almeida, Tahiana Machado Antunes Rodrigues, Isabella Tabelli Bragança, Aline Alves Scarpellini Campos, Verônica Baú Manzoni, Lais Ceschini Machado, Luisa Maria Inácio da Silva, André Luiz Sá de Oliveira, Marcelo Henrique Santos Paiva, Zenaida Marion Alves Nunes, Paula Rodrigues de Almeida, Meriane Demoliner, Juliana Schons Gularte, Mariana Soares da Silva, Micheli Filippi, Vyctoria Malayhka de Abreu Góes Pereira, Fernando Rosado Spilki, Ana Beatriz Gorini da Veiga, Gabriel Luz Wallau

## Abstract

Chikungunya is a reemerging arthropod-borne virus that has been causing large outbreaks in the Americas. In Brazil, Asian-Caribbean and ECSA genotypes have been detected and lead to large outbreaks in several states since 2014. In Rio Grande do Sul (RS), the southernmost State, the first autochthonous cases were reported in 2016. We employed genome sequencing and epidemiological investigation to characterize the increasing CHIKF burden in RS between 2017– 2021. Distinct lineages of the ECSA genotype were responsible for human infections between 2017–2021. Until 2020, CHIKV introductions were most travel associated and transmission was limited. Then, in 2021, the largest outbreak occurred in the state associated with the introduction of a new ECSA lineage. New CHIKV outbreaks are likely to occur in the near future due to abundant competent vectors and a susceptible population, exposing more than 11 million inhabitants to an increasing infection risk.

## Introduction

Chikungunya virus (CHIKV) was first reported infecting humans in July 1952 on the Makonde Plateau, which is part of today’s Tanzania. The human disease was named chikungunya as a reference to the patient’s debilitating symptoms, meaning “the one that bends up the joints” (1). There are four known CHIKV genotypes in circulation worldwide, which were named after their geographical region of discovery: East-Central-South-African (ECSA) and West African (WA) genotypes that are endemic/epidemic in sub-Saharan Africa and are mostly transmitted in a sylvatic cycle, and the Asian-Caribbean (AC) genotype that is mostly endemic/epidemic in Southeast Asia and Central America (2). In the last decades, the ECSA genotype underwent major geographical expansion, leading to major outbreaks around the globe (3). Three well characterized expansion events and associated human outbreaks are known: I - the Indian Ocean Lineage (IOL) derived from the ECSA genotype caused explosive epidemics in the Indian Ocean islands and Asia between 2005 and 2011 (4,5); II - several outbreaks occurred in the Pacific Islands with further spread to the Caribbean region (6,7); and III - introduction of the ECSA genotype in South America and Brazil, which was associated with major outbreaks (8,9).

In Brazil, the wide distribution of competent mosquito populations (*Aedes aegypti* and *A. albopictus*) and a large naive human population favor CHIKV spread (10). The first CHIKV infections reported in Brazil were travel-related, resulting in limited onward transmission in 2010 (6,11). In 2014, autochthonous transmissions of two genotypes were detected: the AC genotype in the North region and the ECSA genotype in the Northeast region of the country (8). Early studies characterizing these events suggested that both lineages would spread and cause outbreaks country-wide, with a higher transmission potential in the tropical region (8). This hypothesis was later corroborated with several Brazilian states in the North, Northeast, Southeast and Central-West regions experiencing large CHIKV outbreaks; whereas the South region experienced lower CHIKV incidence (12–19).

Rio Grande do Sul (RS), the subtropical southernmost state of Brazil, first detected CHIKV infections in humans in 2014, but they were mostly travel-related (20). The first autochthonous transmissions occurred only in 2016, but the low number of cases suggested limited onward transmission (20). However, since 2017 an increasing number of suspected CHIKV infections (symptoms-based diagnostic) have been reported in RS (21). CHIKV genotype-based differential molecular diagnostic is still lacking, hence virus genotypes responsible for such outbreaks are still unknown.

Here we describe the diagnosis of Chikungunya-suspected cases in RS state between 2017–2021. In addition, we performed CHIKV genome sequencing to characterize the genotype(s) currently circulating in the state and evaluate the source and sink phylogeographic pattern. Our results show that Chikungunya fever (CHIKF) cases in RS were driven solely by the ECSA genotype in the period studied and that CHIKV introductions in the state between 2017–2020 were mostly a result of travel-related events to/from Brazilian states with high incidence of CHIKF. Moreover, we characterized the largest recorded CHIKF outbreak in RS, which occurred in 2021 and was associated with the introduction of a new ECSA lineage in the NorthWest region of the state.

## Material and Methods

### Patients and Samples

This study included the analysis of serum samples from patients showing typical arbovirus symptoms between 2017 and 2021 sent to the Central Laboratory of Public Health of Rio Grande do Sul (LACEN-RS) for Dengue virus (DENV), Zika virus (ZIKV) or CHIKV diagnosis (**Suppl. Mat. 1**).

### CHIKV detection by RT-qPCR

RNA was isolated from samples collected within 8 days of symptoms onset using either KingFisher Flex System (ThermoFisher Scientific) or Extracta 96 (Loccus) following manufacturers’ instructions. CHIKV detection was based on RT-qPCR using specific primer/probe sets for CHIKV nonstructural NSP1 and NSP4 proteins (**Suppl. Tab. 1**), or using commercial kits (Molecular ZDC Bio-Manguinhos or Molecular ZDC I- Zika/Dengue/Chikungunya/IBMP, Brazil), according to manufacturers’ instructions. Samples showing amplification with RT-qPCR cycle threshold (Ct) ≤38 were considered CHIKV-positive; positive and negative controls were used in each analysis.

### CHIKV antibody tests (IgM/IgG/MAC-ELISA)

Samples from patients with suspected CHIKV infection collected between 9 and 30 days of symptoms onset were analyzed using anti-CHIKV IgM ELISA kit (Euroimmun, Germany; Dia.Pro, Italy; or Vircell, Spain) according to manufacturers’ instructions. IgM-reactive samples were confirmed by CHIKV-specific IgM capture enzyme-linked immunosorbent assay (MAC-ELISA, Euroimmun, Germany) in 2018–2019 according to manufacturer’s instructions. Samples collected after 30 days of symptoms onset were tested using anti-CHIKV IgG ELISA kit (Euroimmun, Germany or XGEN/XG-CVG- MB, Brazil) following manufacturers’ instructions (**Supplementary Table 1**). All cases that were either IgM or IgG-reactive and showed clinical and epidemiological CHIKV infection characteristics were considered CHIKV-positive.

### Ethical aspects

The use of anonymized samples and metadata from patients was approved by the Ethical Committees of Aggeu Magalhaes (CAAE 10117119.6.0000.5190), Universidade Federal de Ciências da Saúde de Porto Alegre (CAAE 43338715.8.0000.5345), and LACEN-RS (n. 371.278).

### Spatial distribution

CHIKV-confirmed cases spatial distribution per municipality was added to the digital cartographic base containing the municipalities of RS state, in shapefile format (shp). Latitude/Longitude Geographic Projection System and the Geodetic Reference System SIRGAS 2000 were collected from the Brazilian Institute of Geography and Statistics (IBGE) website. Due to the large differences in the numbers of CHIKV cases registered per year, we transformed the raw number of cases by the logarithm function in base 10. To generate the spatial surface, the spatial interpolation method Inverse Distance Weighted (IDW)(22) was used. The period of the study was divided by year to verify the spatial dynamics of the cases over time. The program QGIS 3.10 (Open Source Geospatial Foundation) was used for data insertion, spatial analysis and map generation.

### Genome sequencing

We conducted genome-wide amplification and sequencing of all samples that had a positive CHIKV RT-qPCR (Ct ≤ 30). Amplification was performed with a primer set described by Machado et al. 2019 (23). Libraries were prepared using either the Nextera DNA Flex Library Preparation Kit (Illumina Inc.), or an adaptation of the COVIDseq protocol (https://www.illumina.com/products/by-type/ivd-products/covidseq.html) where SARS-CoV-2 primers were replaced by CHIKV primers sets. Sequencing was performed using Illumina MiSeq platform.

### Genome assembly and phylogenetic analysis

A reference-based genome assembly was conducted using the ViralFlow v.0.0.6 pipeline (24). The closest and most well annotated reference CHIKV genome KP164568.1 was chosen for read mapping. Consensus genomes were obtained by majority rule considering a valid base that showed coverage depth ≥5. Genomes that showed >95% of coverage breadth were further selected for phylogenetic analysis. To genotype and characterize the transmission chains of the CHIKV lineages, we downloaded all near complete CHIKV genomes (≥9000bp) from NCBI (https://www.ncbi.nlm.nih.gov/) and ViPR (25) databases, and removed redundancy. Then we built two datasets for phylogenetic reconstruction: I - a full dataset totalling 1,407 genomes, including genomes from the three known CHIKV genotypes; II - a second dataset based on the maximum likelihood phylogenetic analysis of the full dataset employing IQ-TREE 2.1.2 (26). Based on monophyletic clades (aLRT ≥ 80), we extracted the second dataset covering all genomes sequenced from Brazilian states and South American countries (255 genomes). We kept only genomes containing associated geographical (South America country except Brazil and Brazilian states) and sampling date information. We used Mafft v7 (27,28) and Aliview 1.28 (29) for alignment and visualization, and TempEst v.1.5.3 (30) to evaluate the temporal signal of the dataset. A Bayesian phylogenetic approach was used to reconstruct the evolutionary relationships using Beast 1.10.4 (31), and a discrete phylogeographic analysis was performed using Brazilian states and South American countries sampling collections as discrete characters (32). Since travel information was available for some samples from RS, we adjusted the xml files with this information to inform the source and sink discrete location following (https://beast.community/travel_history). Sampling and mixing of posteriors were accessed using Tracer 1.7.2 (33). The effective sampling size ≥200 was used as a minimum threshold for all parameters estimated. We used 500 million chains sampling every 10,000 and a burnin of 10% using TreeAnnotator. A relaxed molecular clock and a bayesian coalescent skyline prior were used for tree reconstruction and a strict clock prior for discrete state jumps.

## Results

### CHIKV cases in RS between 2017–2021

Chikungunya-confirmed cases in Rio Grande do Sul state varied from 14 to 52 between 2017-2020 (52 in 2017, 42 in 2018, 33 in 2019, and 14 in 2020) (**Figure 1A**). Several patients reported prior traveling to other Brazilian states with known endemic CHIKV transmission before symptoms onset (10 in 2017, 6 in 2018, 6 in 2019, 3 in 2020), such as Rio de Janeiro (Rio de Janeiro State), Fortaleza (Ceara State) and other Brazilian states including Bahia, Minas Gerais and Pernambuco (**Supplementary Material 1**). The geographical distribution of cases comprehended distinct regions of RS, with the number of cases in each region varying along the period, except for the Porto Alegre metropolitan region that consistently reported confirmed cases (**Figure 1A**).

**Figure 1.**
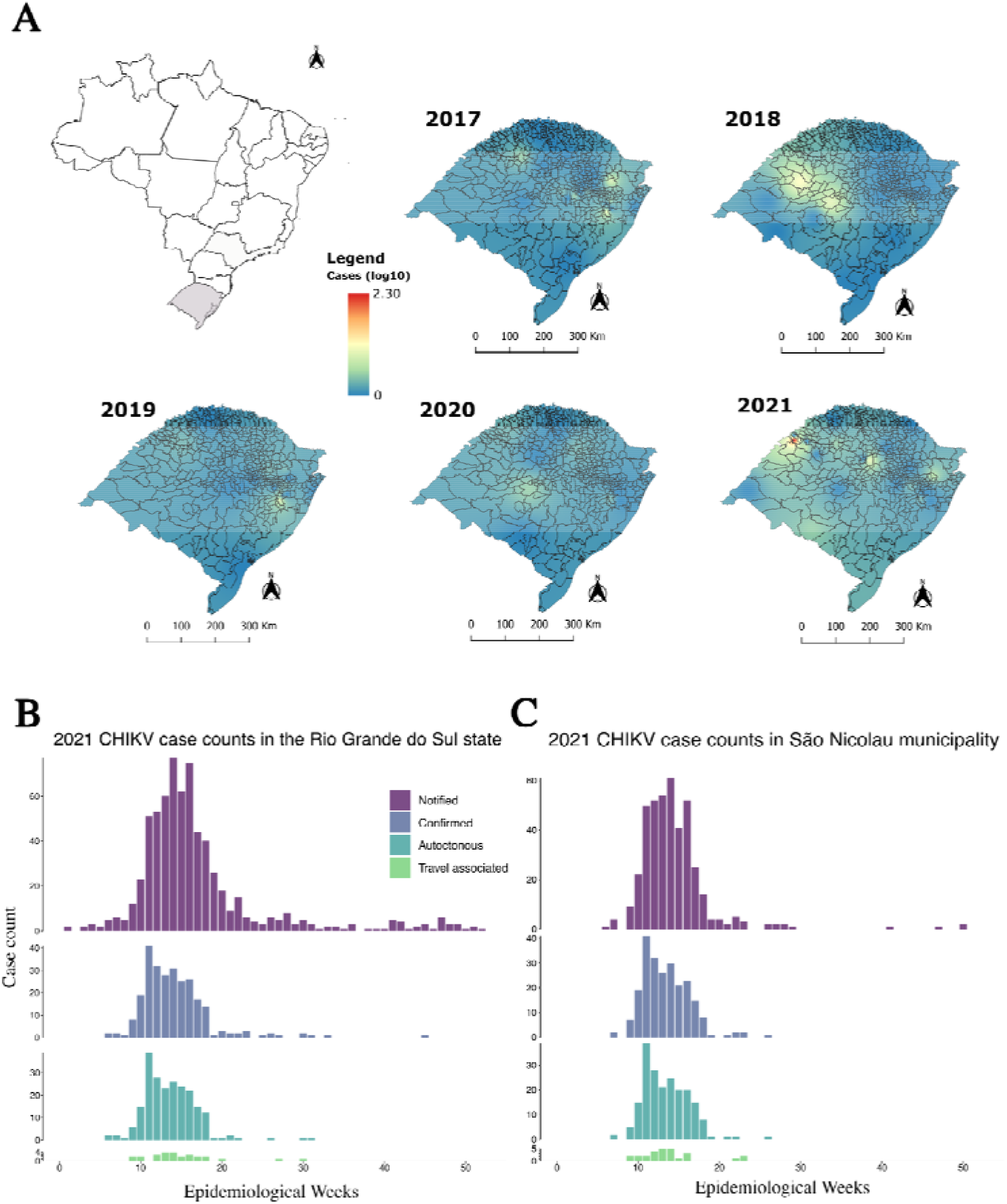
Spatial interpolation of CHIKV cases in the state of Rio Grande do Sul, Brazil, between 2017 and 2021transformed to logarithms to the base 10 (**A**). Graphs show the number of CHIKV cases notified (suspected based on typical CHIKF symptoms), confirmed by molecular assays (RT-qPCR or Immuno assays), showing autochthonous transmission (no travel- associated) and confirmed cases with travel to/from endemic Latin American or Brazilian regions prior to symptoms onset in Rio Grande do Sul state (**B**) and São Nicolau municipality (**C**).

In 2021, besides sporadic confirmed cases in east, central and south regions of the state, a CHIKV outbreak occurred in São Nicolau, a small municipality in the Northwest region of the state with around 5,700 inhabitants (**Figure 1A**). This outbreak was characterized by 220 confirmed cases, surpassing the total of cases observed in the previous four years in RS (141 confirmed cases), being the largest outbreak ever reported in the state (**Figures 1A and B)**. The second municipality with the highest number of cases in RS in 2021 was Espumoso, with only 10 cases (**Figure 1B and Supplementary Material 1**). In the following section we describe the São Nicolau outbreak in detail.

### The 2021 São Nicolau outbreak

A total of 9,309 samples of patients from São Nicolau with suspected arboviral disease in 2021 were analyzed for investigation of DENV, ZIKV, and CHIKV. Of these, 415 presented typical CHIKF symptoms (acute fever, joint and muscle pain, headache, nausea, fatigue, exanthema, and severe joint pain) (34,35), 220 of which (53.%) were confirmed as CHIKV-positive using either RT-qPCR (84, 36.7%) or Immunological-based assays IgM and/or IgG (136, 61.8%). Epidemiological investigation showed that 202 (91.8%) of the confirmed cases likely derived from autochthonous acquired infection with no travel history to other Brazilian endemic states or South American countries, while 18 (8.2%) reported travel previous to symptoms onset. Of the remaining 195 CHIKV negative patients, six were positive for DENV and four for ZIKV, while the remaining were negative for all arboviruses investigated.

The first identified case of CHIKV infection in Sao Nicolau was confirmed by RT-qPCR in a sample collected on March 17^th^ (Epidemiological Week - EW 11), from a 52 years old healthcare worker who presented intense arthralgia and exanthem symptoms on March 9^th^ (EW 10). Following this first detection other patients presenting fever, arthralgia, myalgia, joint swelling, retro-orbital pain, and/or exanthema were investigated and confirmed CHIKV infection or exposure in March 2021 (EW 9–13). Then a retrospective investigation of arbovirus-suspected cases was conducted, confirming the CHIKV outbreak in the municipality. The outbreak occurred from February to July 2021 (EW 7–26), with the peak in March/April (EW 9–17) (**Figure 1B and C**). Travel-associated infections were also notified between EW 9–23, and included individuals who had been in different endemic Brazilian states (**Figure 1C**). Most reported symptoms (>10%) of CHIKF-confirmed cases were arthralgia, fever, headache, rash, myalgia, nausea, back pain and vomiting (**Figure 2A**). No patient required hospitalization and no fatalities were reported. Most patients with CHIKV infection (76.2%) were adults between 30 and 79 years of age (**Figure 2B**).

**Figure 2.**
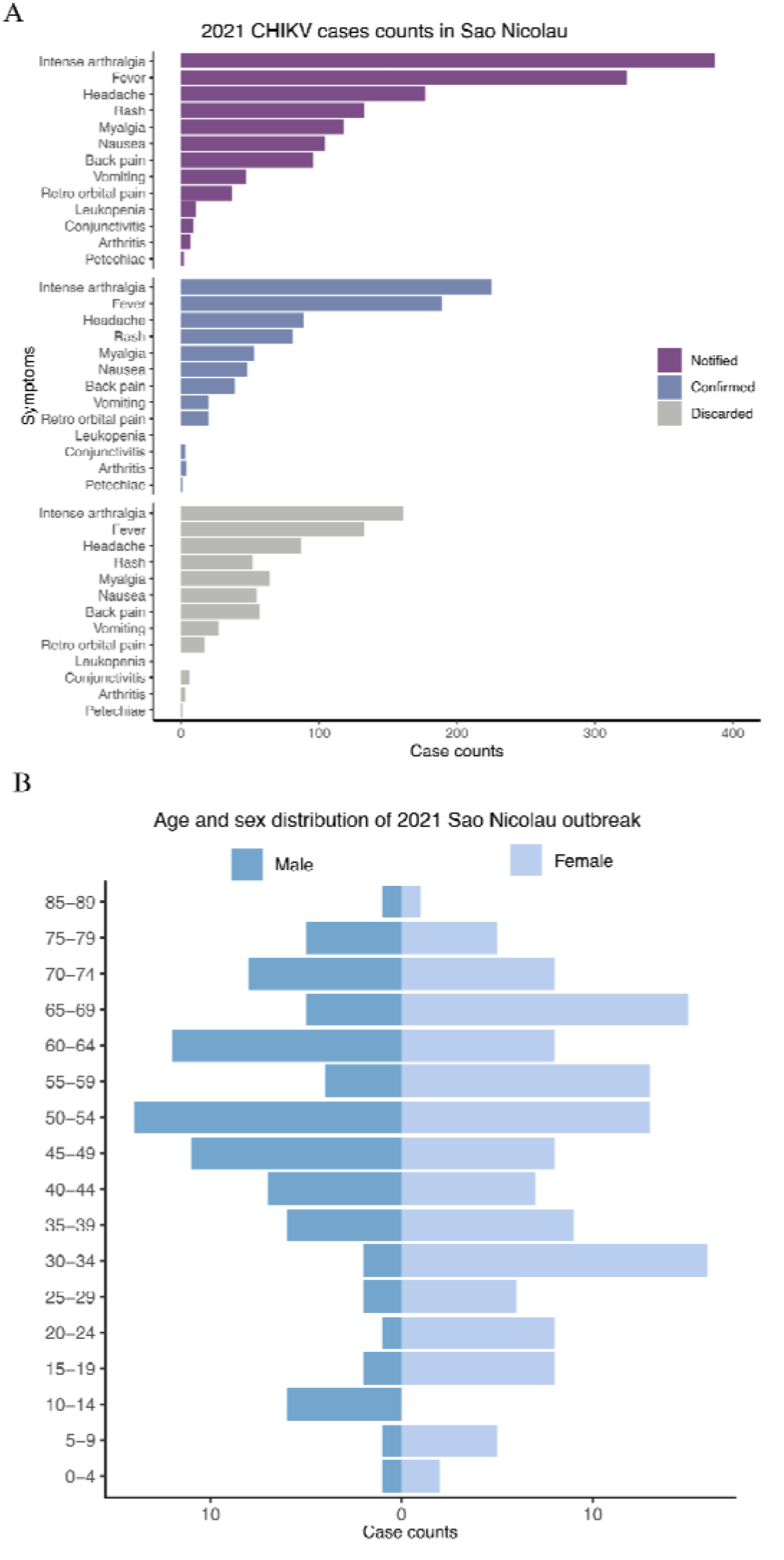
Symptoms and age distribution of cases with suspected, confirmed and discarded CHIKV infection. (**A**) Number of CHIKV cases notified (suspected based on typical CHIKF symptoms), confirmed by molecular assays (RT-qPCR or Immuno assays) and discarded from São Nicolau state in 2021 stratified by reported symptoms; and (**B**) age stratification of cases in São Nicolau municipality.

To characterize and investigate in more detail cases suspected of CHIKV infection with a negative CHIKV RT-qPCR, we plotted the frequency of symptoms, for notified, confirmed and discarded cases from São Nicolau municipality (**Figure 2A**). No differences in the symptoms presented by negative and positive patients were observed (**Figure 2**).

### CHIKV whole-genome sequencing and phylogenetic analysis

We sequenced 23 near-complete CHIKV genomes (one sampled in 2017, four in 2019 and 18 in 2021) (**Table 1**), showing average coverage breadth and depth or 98.5 and 3,071 respectively (**Table 1**).

**Table 1.** Information of samples and genomes sequenced from Rio Grande do Sul and São Nicolau outbreak between 2017–202.

| Sample ID | Municipality | Symptoms started (dd-mm-yyy) | Sampling date (dd-mm-yyy) | Travel association | Ct value | Average Depth | Coverage Breadth |
| --- | --- | --- | --- | --- | --- | --- | --- |
| 1115 | São Nicolau | 3/9/2021 | 3/17/2021 |  | 24 | 4741.98 | 97.55 |
| 1124 | São Nicolau | 3/13/2021 | 3/17/2021 |  | 31 | 3629.10 | 99.22 |
| 1499 | São Nicolau | 3/18/2021 | 3/24/2021 |  | 24 | 5444.54 | 99.26 |
| 1502 | São Nicolau | 3/18/2021 | 3/24/2021 |  | 30 | 2557.28 | 100.00 |
| 1504 | São Nicolau | 3/15/2021 | 3/24/2021 |  | 30 | 4454.54 | 99.46 |
| 1509 | São Nicolau | 3/21/2021 | 3/24/2021 |  | 21 | 5623.90 | 99.91 |
| 1892 | São Nicolau | 3/29/2021 | 3/30/2021 |  | 20 | 3773.15 | 99.75 |
| 2405 | São Nicolau | 3/26/2021 | 3/31/2021 |  | 30 | 2053.71 | 96.86 |
| 4079 | São Nicolau | 4/13/2021 | 4/15/2021 |  | 25 | 3383.81 | 99.89 |
| 4080 | São Nicolau | 4/9/2021 | 4/16/2021 |  | 29 | 1273.39 | 98.97 |
| 4081 | São Nicolau | 4/15/2021 | 4/15/2021 |  | 28 | 5643.75 | 99.99 |
| 5166 | São Nicolau | 4/19/2021 | 4/23/2021 |  | 26 | 2599.88 | 99.89 |
| 5171 | São Nicolau | 4/16/2021 | 4/23/2021 |  | 28 | 3711.22 | 99.91 |
| 5180 | São Nicolau | 4/19/2021 | 4/26/2021 |  | 28 | 4601.83 | 98.54 |
| 5183 | São Nicolau | 4/21/2021 | 4/27/2021 |  | 30 | 2162.92 | 97.09 |
| 6165 | Espumoso | 5/4/2021 | 5/7/2021 |  | 28 | 2214.65 | 97.21 |
| 6701 | Espumoso | 5/8/2021 | 5/12/2021 |  | 28 | 1289.84 | 97.04 |
| 7302 | São Nicolau | 5/10/2021 | 5/13/2021 |  | 30 | 1258.46 | 95.76 |
| 2849 | Canoas | 6/19/2019 | 6/19/2019 |  | 21 | 2900.56 | 97.84 |
| 1982 | Rio Grande | 5/16/2019 | 5/18/2019 | Rio de Janeiro | 35 | 1955.66 | 98.58 |
| 158 | Porto Alegre | 1/28/2019 | 2/2/2019 | Cuba | 35 | 2072.16 | 98.56 |
| 336 | Porto Alegre | 3/6/2019 | 3/9/2019 |  | 23 | 198.69 | 97.41 |
| 1328 | Caxias do Sul | 4/26/2017 | 5/3/2017 |  | 18 | 3088.81 | 98.70 |

Whole genome phylogenetic reconstruction of all CHIKV genomes available recovered the three major genotypes: WA, ECSA and AC (**Supplementary Figure 1A**). All genomes obtained in this study clustered with high branch support (ultrafast bootstrap 97) within the ECSA genotype (**Supplementary Figure 1A and B**). In addition, we investigated in more depth the phylogenetic clustering of the RS CHIKV genomes with all genomes from other Brazilian states and South American countries. Phylogenetic reconstruction was performed using the smaller database II (**see Material and Methods and Supplementary Figure 1B**). RS genomes clustered in four highly supported clusters (1–4 in **Suppl. Fig. 1B, Suppl. File 1**). Cluster 1 grouped a RS genome from 2017 with three genomes from Ceara and Rio de Janeiro states sampled in 2017 and 2019, respectively. Cluster 2 grouped a 2019 RS genome within a clade comprising several genomes from Mato Grosso, Para, Maranhao states, and a single genome from Amazonas state. Clade 3 comprises a large clade with several samples from Rio de Janeiro and two 2019 genomes from RS; and Clade 4 included one RS genome from 2019, all samples from São Nicolau 2021 outbreak, and a sister clade comprising of several genomes from Sao Paulo state (**Supplementary Figure 1B**).

Molecular clock phylogenetic analysis estimated that clusters 1 and 2 last common ancestor (LCA) occurred in April 2016 (HPD 95 = January/June 2016) and May 2016 (HPD 95 = January/October 2016) (**Suppl. Mat. 2**). Clade 3 LCA was dated from December 2017 (HPD 95 = October 2017/February 2018), while the Clade 4 LCA occurred around November 2018 (HPD 95 = June 2018/May 2019) (**Supplementary Material 2**). The lineage leading to the São Nicolau outbreak was likely introduced in the state around October 2020 (HPD 95 = June 2020/January 2021) and remained cryptically circulating in the region until the first cases in February 2021 (**Figure 1C**).

### Discrete phylogeographic analysis

Bayesian discrete phylogeographic analysis detected four introductions of CHIKV in RS. A single 2017 genome shared a LCA that was likely circulating in Bahia state (ASR PP of 0.73, Clade 1 - **Supplementary Figure 1B**). Four 2019 genomes originated likely from three independent introductions: Clade 2 shared a LCA in the Para state (ASR PP of 0.59), Clade 3 and 4 shared a LCA from Rio de Janeiro state (PP = 1 and 0.86). The two 2019 genomes forming Clade 3 were recovered from RS patients with different travel histories: while one reported no travel history, the second (patient 158) had traveled to Cuba before the symptoms onset (**Table 1**). The discrete phylogeographic analysis reconstructed the ancestral of these genomes as originating in RS, but with a low state reconstruction support (ASR PP 0.40). Therefore, patient 158 either has acquired CHIKV infection while traveling in Cuba or was infected in RS after arrival. The second hypothesis is more likely, since the ECSA genotype is more widespread in Brazil and South America, while the AC genotype was more predominant in Central America and Caribbean islands (36–38), including Cuba (39). Lastly, Clade 4 included a large cluster of genomes from the São Nicolau 2021 outbreak that clustered with several genomes from Sao Paulo state and with one RS genome from 2019 (patient 1982) in a basal position (**Supplementary Figure 1B**). Patient 1982 resided in RS, but had traveled to Rio de Janeiro before symptoms onset. Therefore, after incorporating the travel history of this patient, the common ancestor of Clade 4 was more likely circulating in Rio de Janeiro (ASR PP = 0.86) (**Figure 3**).

**Figure 3.**
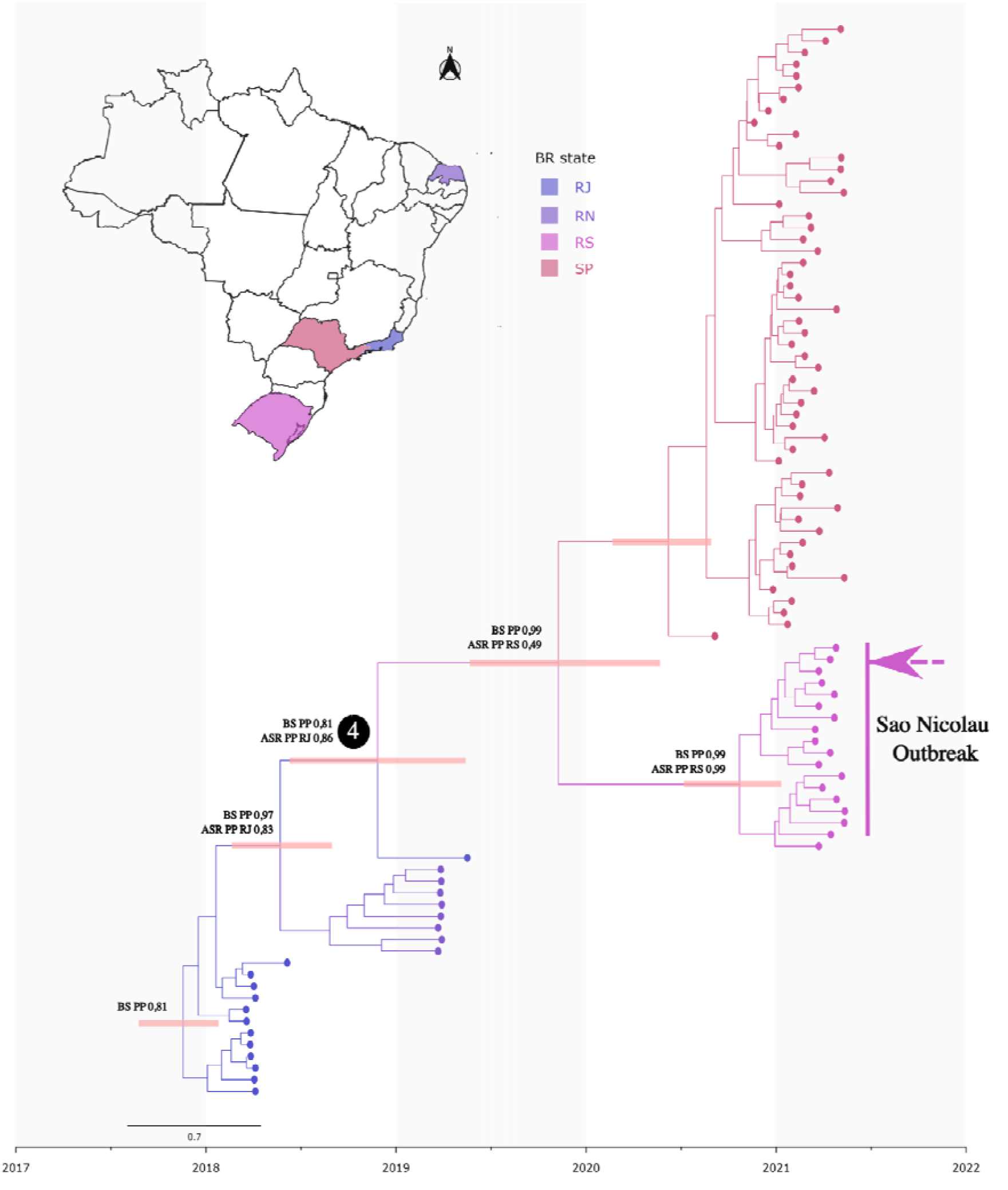
Time resolved phylogenetic tree reconstructed using a bayesian phylogenetic approach and Brazil map showing Brazilian states sampling location of CHIKV genomes included in the analysis. BS PP:Bayesian Support Posterior Probability; ASR PP: Ancestral State Reconstruction Posterior Probability.

## Discussion

Chinkungunya is an arbovirus, transmitted to humans by mosquitoes of the *Aedes* genus, *A. aegypti* and *A. albopictus* (40). Most CHIKF cases show symptoms associated with medium to severe morbidity; neurological complications and patient death may also occur (35,41). Both AC and ECSA lineages are expanding their range, causing large human outbreaks in the Americas, Europe and Southeast Asia (6,42). In Brazil, the AC and the ECSA lineages were introduced in 2014 and simultaneously spread in the country. However, after the first years of expansion, these genotypes have been undergoing distinct transmission dynamics. While the AC genotype remained mostly restricted to the North region (13), the ECSA genotype is spreading widely and has been detected in all Brazilian states (9,12,14,15,17,17–19). A recent study showed that the most affected region of Brazil was the Northeast (16), whereas limited CHIKF cases had been detected in the South region (20,43). In our study, we found that different ECSA lineages were introduced in RS, the southernmost Brazilian state, between 2017–2020, and that the São Nicolau 2021 outbreak was caused with the arrival of a new lineage, which was circulating four months previous to the detection of the first CHIKF case in that municipality.

*A. aegypti*, the main vector associated with CHIKV transmission in Brazil, is widely distributed across the country, including RS (21), and increasing human infection with distinct arboviruses (i.e DENV, ZIKV, CHIKV) have been reported (44). Arboviruses infections in RS follow a broad seasonal epidemic variation as characterized for DENV in Brazil, where most cases occur between April and June (43). The RS state presents distinct climatic seasons, hence human cases of arboviral infections are mostly restricted to the hot and humid summer in the state (20). Nonetheless, during the 2021 São Nicolau outbreak, cases occurred until July 2021, even with the lower winter temperatures (average of 14.8°C) that are expected to reduce vector population and virus transmission. It is important to note that São Nicolau reported its first arbovirus infection in 2021, and the following CHIKV outbreak was the largest recorded in the state. Moreover, based on symptom stratification of the negative 195 samples, it is likely that a large proportion are derived from the same CHIKV outbreak, since only 10 samples were positive for ZIKV and DENV virus. However, it remains to be assessed if other unknown or not assessed arboviruses may have been cocirculating. Lastly, we detected that the majority of the cases are autochthonous cases, but travel-associated cases were also identified. Taken together, the most likely scenario to explain the São Nicolau outbreak is that in 2021 a new ECSA lineage introduction likely derived from a travel-associated importation established extensive communitary transmission through competent vector mosquitoes.

It is important to note that this study has some limitations. Individuals spontaneously sought municipal health care units, and many samples were collected after 9 days of symptoms onset; therefore, a large fraction of samples were only accessed by immunoassay methods due the lower probability of detecting viral RNA from samples. This not only decreased diagnosis sensibility but also hindered genomic analysis of all samples; nonetheless, the CHIKV genomes obtained in this study are likely to represent the CHIKV lineages circulating in RS. Noteworthy, there is no systematic CHIKV genomic surveillance in Brazil, which may have impacted the phylogeography analysis. Therefore, our results should be interpreted with caution and further evaluation of the phylogenetic and phylogeography hypothesis raised in this study is needed.

Climatic conditions in subtropical regions of the globe such as the RS state are changing substantially in the last years, with longer and hotter summers (45), providing a wider transmission season for arboviral diseases. Approximately 1.3 billion people live in high risk areas for CHIKV infection around the globe (46,47), and climate modeling studies suggest that previously unaffected areas may now provide suitable conditions for arbovirus transmission (46). The association of competent vector mosquitoes, the large susceptible human population (no antibody barrier) in RS and the lack of pharmacological treatment options point to a harsh scenario of increased infection risk for the 11 million inhabitants of the state. Once CHIKV infection induce life-long immunity (48), it is likely that following outbreaks may happen in other RS municipalities, but it remains to be assessed if new outbreaks will be triggered by previous lineages overwintering in the state or due to continuous introduction of new lineages from other endemic regions in the years to come.

The increasing arbovirus burden in subtropical and temperate regions is in line with more broad analysis of increasing risk of arthropod-borne infections in subtropical and temperate regions of the globe due to environmental and climate changes, including global warming (49,50). Therefore, the countries and states of the Southern Cone region of South America should better prepare implementing early warning systems, better virus-vector surveillance, control practices and population educative measures to cope with the increasing arboviruses’ impact in the future.

## Acknowledgements

This work was supported by the Health Department of Rio Grande do Sul and Brazilian Ministry of Health. ABGV and GLW hold fellowships from Conselho Nacional de Desenvolvimento Científico e Tecnológico (Grant processes 306369/2019-2 and 303902/2019172-1, respectively).

## Data Availability

Genome sequences have been deposited in GenBank under accession numbers (will be made available upon acceptance of the publication).

